# ReCycled: A Tool to Reset the Start of Circular Bacterial Chromosomes

**DOI:** 10.1101/2025.04.07.647662

**Authors:** Vincent Somerville, Michael Schmid, Matthias Dreier, Philipp Engel

## Abstract

**Summary (two sentences):** For many publicly available bacterial genomes the chromosome start is not set at the replication initiation protein. This is a burden for comparative genomics studies as the synteny between such genomes cannot readily be assessed. Here, we present ReCycled, a tool for identifying and resetting the start of circular bacterial chromosomes.

**Availability and implementation:** Freely available on GitHub at https://github.com/Freevini/ReCycled under the GPL-3.0 license. Runs on all tested GNU/Linux systems.

**Contact:** Vincent Somerville

**Supplementary Information:** The analyses, scripts and data to test the different pipelines are deposited online on zenodo (10.5281/zenodo.15170502).

## Introduction

The number of completely assembled, circular bacterial chromosomes has steadily increased in recent years due to the affordability of long read sequencing technologies. Complete genome assemblies facilitate the comparison of important genomic features of bacteria, such as the conservation of the overall genetic organisation (genome synteny) (Fig. 1A). Most bacterial chromosomes are circular. Hence, an important prerequisite for comparing the overall genomic structure across genomes is that the start of the linearized contig representing the complete bacterial chromosome is set to the same position. Common practice is to set position 1 of the bacterial chromosome (close) to the first position of the replication initiator protein *dnaA* (Hunt et al. 2015). Although there can be exceptions, the replication initiator protein dnaA usually lies close to the origin of replication (OriC) (Sernova and Gelfand 2008).

**Figure 1.**
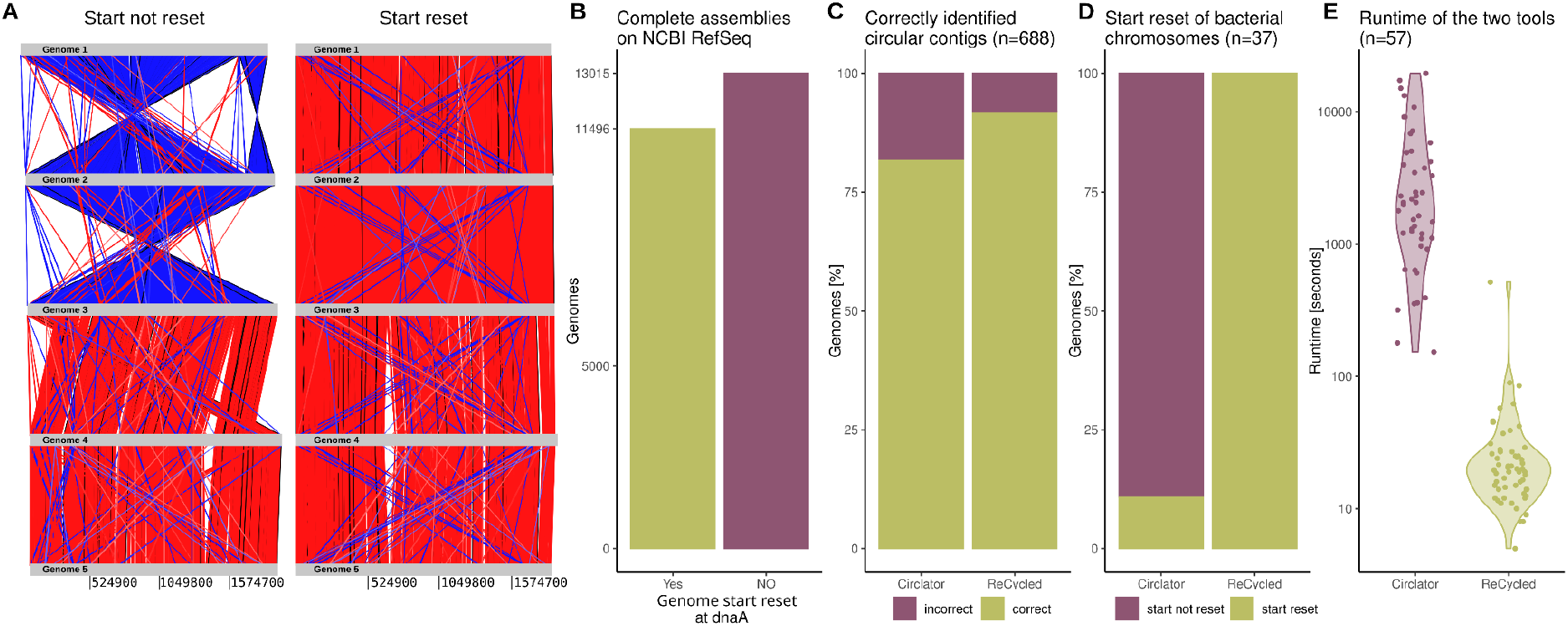
Utility of ReCycled and its performance compared to Circulator. A) Genome synteny plot of five complete *Streptococcus thermophilus* genomes from (Somerville et al. 2021) before and after resetting the start position with Recycled. The plots were created with Artemis (Carver et al. 2008). Red indicates genomic blocks of the same orientation, while blue are blocks of reverse orientation. B) The majority of complete genomes on NCBI RefSeq are not start-aligned, i.e. the replication initiation protein gene *dnaA* is not located in close proximity to the chromosome start. C) The ability to correctly identify circular contigs with Circlator and ReCycled. D) The ability to correctly reset the start of 36 randomly picked non-startaligned but circular RefSeq genomes with Circlator and ReCycled. E) The runtime to circularise genomes with Circlator and ReCycled.

Currently, there are 54,601 complete (and circular) genomes available on the NCBI bacterial RefSeq (April 2025). However for 52% of the chromosomes, the start of the chromosome is not set to the replication initiation protein (Fig. 1B). This can be explained by the fact that only a few genome assembly tools, e.g. Unicycler (Wick et al. 2017) and Trycycler (Wick et al. 2021), set the start of the chromosome in their workflow to a specific position. Several standalone tools are available for identifying the origin of replication or reorienting bacterial genomes, but each comes with specific limitations. Ori-Finder is a web-based tool that predicts the origin of replication; however, it does not reset the genome start point (Gao and Zhang 2008). SeqKit offers a restart command that reorients contigs, but it neither verifies circularity nor detects replication initiation proteins (Shen et al. 2016). Dnaapler, a more recent tool designed for circularizing bacterial chromosomes, also lacks functionality to determine whether a contig is truly circular (Bouras et al. 2024). Circlator is the most comprehensive toolkit that detects circular genomes and resets the chromosomal start position (Hunt et al. 2015). However, it applies a stringent and computationally intensive workflow involving local reassembly and polishing of long-read data. As a result, Circlator is not well-suited for high-throughput applications or for processing large genome datasets in parallel.

## ReCycled: application and usage

Here we present ReCycled: a simple, accurate and fast tool that identifies, circularises and restarts circular bacterial chromosomes. This tool was developed within the framework of a student course in 2020 and has been used and updated since then (Somerville et al. 2025). The workflow of ReCycled comprises five steps. First, the correct installation of the tool (and its dependencies) is verified, the input files are checked for the correct format and the individual genomic contigs are split to separate FASTA files. Second, the replication initiator protein is located by searching against a custom-built replication initiator protein database. The reference database is created by combining all identified replication initiation proteins from the DoriC reference database (Luo and Gao 2018), from Unicycler (Wick et al. 2017) and from the complete genomes of the RefSeq database (29th Dec 2021). Third, the circularity is inferred by checking for i) overlapping contig edges, ii) contig-edge spanning long reads and iii) contig-edge spanning short reads. Only the ii) option is mandatory to be fulfilled. Fourth, position 1 of the circular contigs are reset to ten bases upstream of the identified gene encoding the replication initiator protein. Fifth, the correctness of the restart is verified by locating the replication initiation gene, the contigs are concatenated back together and a log file summarizing the results of the analysis is produced. The workflow is built on a minimal set of default Unix commands as well as Minimap2 (Li 2021), Bedtools (Quinlan 2014) and SeqKit (Shen et al. 2016).

## The performance of ReCycled

In order to check the performance of ReCycled we randomly selected and downloaded the long read data of 52 randomly selected non-start-aligned NCBI RefSeq genomes from the NCBI Short Read Archive (SRA) (Fig. 1B). The long reads were assembled with the *de novo* genome assembler Flye (Kolmogorov et al. 2019). Based on the inspection of the Flye assembly graph, we identified a total of 37 circular bacterial chromosomes in the 52 RefSeq genomes. Moreover, in total 688 circular and non-circular contigs were assembled. We ran ReCycled and Circlator (Hunt et al. 2015) to check the ability to identify circularity and to reset the start position of the circular bacterial chromosomes.

We first checked the ability of the two programs to correctly identify circular contigs. ReCycled was able to correctly predict the circularity of 633 of the 688 (92%) contigs (Fig. 1C). The few contigs missed by ReCycled contained contig edges with repeats or were of low coverage. Importantly, none of the non-circular bacterial chromosomes was incorrectly identified as being circular. Circulator correctly predicted the circularity in 564 of the 688 (82%) contigs.

We next checked for the ability of the two programs to correctly reset the start position of the circular bacterial chromosomes. ReCycled was able to reset the start of all 37 bacterial chromosomes that were assembled and flagged as circular by the *de novo* genome assembler Flye (Kolmogorov et al. 2019) (Fig. 1D). Circlator managed to reset the start of 4 of the 37 (11%) circular bacterial chromosomes.

Finally, we looked at the runtime of the two programs. ReCycled took on average 33 seconds per genome (Fig. 1E) to run, while Circulator took on average 3854 seconds. The difference in runtime can be explained by the fact that Recycled only includes the minimal steps to reset the start position of the bacterial chromosome. In contrast to Circulator, ReCycled neither assembles the contigs into scaffolds nor does it polish the contigs after restarting. Further, the current version does not handle fragmented short read assemblies.

## Conclusions

ReCycled is a reliable, accurate and fast tool to reset the start position of circular bacterial chromosomes (Fig. 1). It contains only a few dependencies and can easily be integrated into automated genome analysis workflows. It runs on Linux based operating systems and therefore can be executed on high-performance computing environments. The tool is freely available under the GPL-3.0 license on GitHub: https://github.com/Freevini/ReCycled.

## Acknowledgments

We thank Aiswarya Prasad and Aline Cuénod for testing the tool and reading the manuscript. This work was supported by the following grants: NCCR, SNF, ERC, Agroscope.

